# Dynamic monitoring of viral gene expression reveals rapid antiviral effects of CD8 T cells recognizing the HCMV-pp65 antigen

**DOI:** 10.1101/2023.07.11.548645

**Authors:** Fawad Khan, Thomas R. Müller, Bahram Kasmapour, Mario Alberto Ynga-Durand, Britta Eiz-Vesper, Jens von Einem, Dirk H. Busch, Luka Cicin-Sain

**Author notes:** These authors contributed equally (Fawad Khan, Thomas R. Müller).

## Abstract

Human Cytomegalovirus (HCMV) is a betaherpesvirus that causes severe disease in immunocompromised transplant recipients. Immunotherapy with CD8 T cells specific for HCMV antigens presented on HLA class-I molecules is explored as strategy for long-term relief to such patients, but the antiviral effectiveness of T cell preparations cannot be efficiently predicted by available methods. Therefore, we developed an Assay for Rapid Measurement of Antiviral T-cell Activity (ARMATA) by real-time automated fluorescent microscopy and used it to study the ability of CD8 T cells to neutralize HCMV and control its spread. As a proof of principle, we used TCR-transgenic T cells specific for the immunodominant HLA-A02-restricted tegumental phosphoprotein pp65. pp65 expression follows an early/late kinetic, but it is not clear at which stage of the virus cycle it acts as an antigen. We measured control of HCMV infection by T cells as early as 6 hours post infection (hpi). The timing of the antigen recognition indicated that it occurred before the late phase of the virus cycle, but also that virion-associated pp65 was not recognized during virus entry into cells. Monitoring of pp65 gene expression dynamics by reporter fluorescent genes revealed that pp65 was detectable as early as 6 hpi, and that a second and much larger bout of expression occurs in the late phase of the virus cycle by 48 hpi. Since transgenic (Tg)-pp65 specific CD8 T cells were activated even when DNA replication was blocked, our data argue that pp65 acts as an early virus gene for immunological purposes. Therefore, ARMATA does not only allow same-day identification of antiviral T-cell activity, but also provides a method to define the timing of antigen recognition in the context of HCMV infection.

## Introduction

Human Cytomegalovirus (HCMV) is a highly common cause of opportunistic infections and disease in immunocompromised transplant recipients (1). HCMV is a member of the beta-herpesvirus subfamily, latently persisting in the host for life upon primary infection, yet able to reactivate its lytic replication cycle, especially in case of immunosuppression (2), and cause life-threatening disease in numerous organ systems (3, 4). In that case, antiviral drugs are used in therapeutic, prophylactic or pre-emptive protocols to counter virus replication and pathogenesis, yet drug-resistant HCMV variants may emerge, resulting in therapy failure and adverse prognosis for the patient (5). Therefore, substantial efforts have been invested in the development of T-cell based immunotherapeutics targeting HCMV antigens (6-12). Recent advances in genetic engineering paved the way to express transgenic antigen-specific T-cell receptors (TCR) and thus to use recombinant T cells as therapeutic agents (13, 14). However, defining which among the available solutions are ideally suited for repressing HCMV replication requires clinical testing, because HCMV species are strictly species-specific and the replication of HCMV cannot be efficiently tested in animal models. Recently, we assessed the antiviral capacity of HCMV-specific CD8 T cells recognizing a defined viral antigen in co-cultures with HLA-matched target cells infected with recombinant HCMV reporters lacking immune evasion genes and expressing fluorescent reporter genes (15). The antiviral capacity was clearly measured and increased when immune evasive genes were not present, providing a preclinical model for the testing of antiviral T cells (15). However, in this study we did not compare various T cell clones to quantify their antiviral activity and identify the best candidates.

In another recent publication, we analyzed the *in vivo* ability of TCR-transgenic CD8 T cells to control CMV, by infecting transgenic mice expressing the human HLA-A02 molecule with a recombinant mouse CMV (MCMV) expressing the A02 restricted antigenic peptide NLVPMVATV (NLV) derived from the human CMV protein pp65. Upon transfer of CD8 T cells recognizing these peptide-HLA (pHLA) complexes, immune responses could be measured, as well as their antiviral activity (13). It remained however unclear if the peptide could be recognized in its native context in the human CMV genome and if it may serve as target for antiviral CD8 T cells in these settings.

The pp65-derived antigenic peptide NLV is a major target for immune recognition by the host immune system (16). However, T-cell responses to this antigen did not correlate with reduced virus levels in transplant recipients (17), raising the question whether its immune recognition would result in virus control. The pp65 is the product of the UL83 gene of HCMV, and it is a major component of the viral tegument, repressing interferon responses immediately upon infection (18) and thus promoting immediate-early gene expression (19). While UL83 mRNA can be detected as early as 5 hours post infection (hpi), its expression is particularly strong in the late phase of the virus replication cycle (20) and hence it is deemed to be an early-late viral gene. Recent studies suggest that UL83 mRNA is transcribed in presence of cycloheximide, arguing that this gene may be expressed with immediate-early kinetics (21). Nevertheless, it remained unclear if the low amounts of pp65 protein present in the early stages of the virus lytic replication cycle can act as antigen, or if T cells recognizing this epitope can act against the virus only in the later stages of the virus cycle.

In this work, we investigated the antiviral activity of CD8 T cells bearing transgenically expressed TCRs recognizing the NLV peptide with different binding avidities to the cognate pHLA complex. Their activity was analyzed at high temporal resolution by means of an Assay for Rapid Measurement of Antiviral T-cell Activity (ARMATA) in human cells infected with reporter HCMV. We show that T cells recognizing pp65 with high avidity of TCR binding deter virus growth more efficiently and we show that the effects of their activation can be observed as early as 6-8 hpi, but not earlier. Moreover, we show that blocking DNA replication does not prevent early pp65 expression and T cell responses to virus-infected cells, arguing that pp65 acts as an early antigen in the virus replication cycle.

## RESULTS

### Characterization of antiviral activity of pp65-specific TCR-transgenic T cells in a dynamic virus reduction assay

To assess the neutralization capacity of T cell populations, previously described TCR clones (13) were used to observe the degree of their viral inhibition in the Assay for Rapid Measurement of Antiviral T-cell Activity (ARMATA). The assay is based on real-time monitoring of virus-infected cells in presence of immune cells, as a rapid assay to measure antiviral activity of defined populations of immune cells (15, 22). Here, we used it for preclinical assessment of the efficacy of transgenic-T cells for adoptive immunotherapy. Briefly, NLV-specific transgenic CD8 T cells were generated by retroviral transduction of primary peripheral blood mononuclear cell (PBMC) and added to MRC-5 cells infected with reporter HCMVs expressing fluorescent proteins (22, 23) at various effector to target (E:T) ratios. Since MRC-5 cells carry HLA-A02 molecules on their surface, they can display the NLV peptide to the TCR-transgenic CD8 T cells. Cells were continuously monitored by live cell imaging to define the virus gene expression over time, while cytokine analysis from co-culture supernatants was performed at 24-48 hours post infection (hpi) as measurement of T cell activation (Fig. 1). To assess the sensitivity and robustness of the assay, HLA-A02–restricted pp65-specific CD8 T cells were used in co-culture with cells infected with a reporter HCMV. For our experiments, we used three previously identified TCRs (24) and measured their binding avidity using the K_off_-rate assay (25). We observed largely different binding avidities between the TCRs, with TCR 5-2 showing the longest and TCR 1-4 the shortest half-life time (t_1/2_) (Fig. 2A, B). These cell lines were purified by magnetic cell separation (Fig S1) and used to assess the sensitivity of ARMATA. Since we used pp65-specific CD8 T cells, we used also a recombinant virus where the GFP gene was fused to the UL83 HCMV gene, which encodes the pp65 protein (26) and promptly observed a reduction of viral signal (Fig 2C). We monitored the antiviral activity of T cells by quantifying the GFP signal at E:T ratios of 1:10 or 1:1 over time. We compared the effects in low multiplicity of infection (MOI) of 0.1 (Fig. 2D, representative figure at 96hpi is shown in panel 2C) or in high MOI settings of 1 (Fig. 2E), and noticed reduced GFP signals in the presence of T cells. In line with expectations, the differences in antiviral activity were observed earlier when the E:T ratio was higher. Importantly, the reduction of virus signal correlated closely with binding avidity of TCRs, with TCR 1-4 showing the least, and TCR 5-2 the highest antiviral activity. The GFP signal was reduced in these co-cultures in comparison with control cells infected in absence of T cells or in co-cultures with mock-transfected PBMCs. At MOI of 0.1 and E:T 1:10, the hierarchy in antiviral activity was noticeable in terms of signal amplitudes and in terms of time kinetics; TCRs that reduced the GFP signal to lower levels acted earlier (Fig. 2D, left panel).

**Figure 1.**
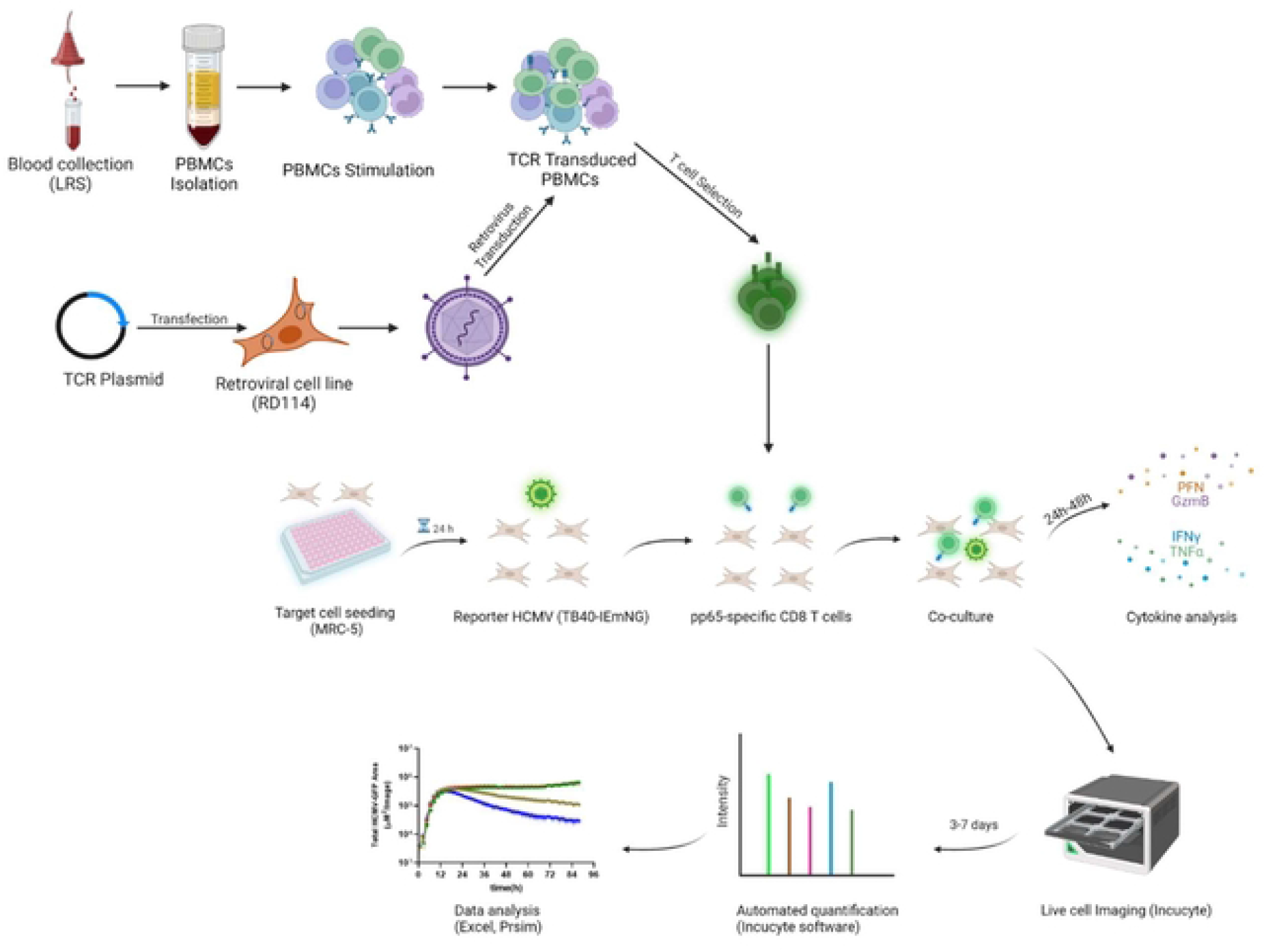
Schematic workflow of the pp65-specific T cells generation and co-culture in ARMATA. PBMCs were isolated and stimulated, followed by retroviral transduction with different TCR Plasmids. For ARMATA, MRC-5 (HLA-A02) cells were seeded and infected with reporter TB40 expressing fluorescent proteins under the control of defined endogenous viral promoters. After infection and washing, pp65-specific CD8 T cells were added to the culture at indicated E:T ratios and co-cultures were monitored by live cell imaging for 3-7 days post infection. Supernatants were collected at indicated times (typically between 24 and 48 hpi) for cytokine analysis.

**Figure 2.**
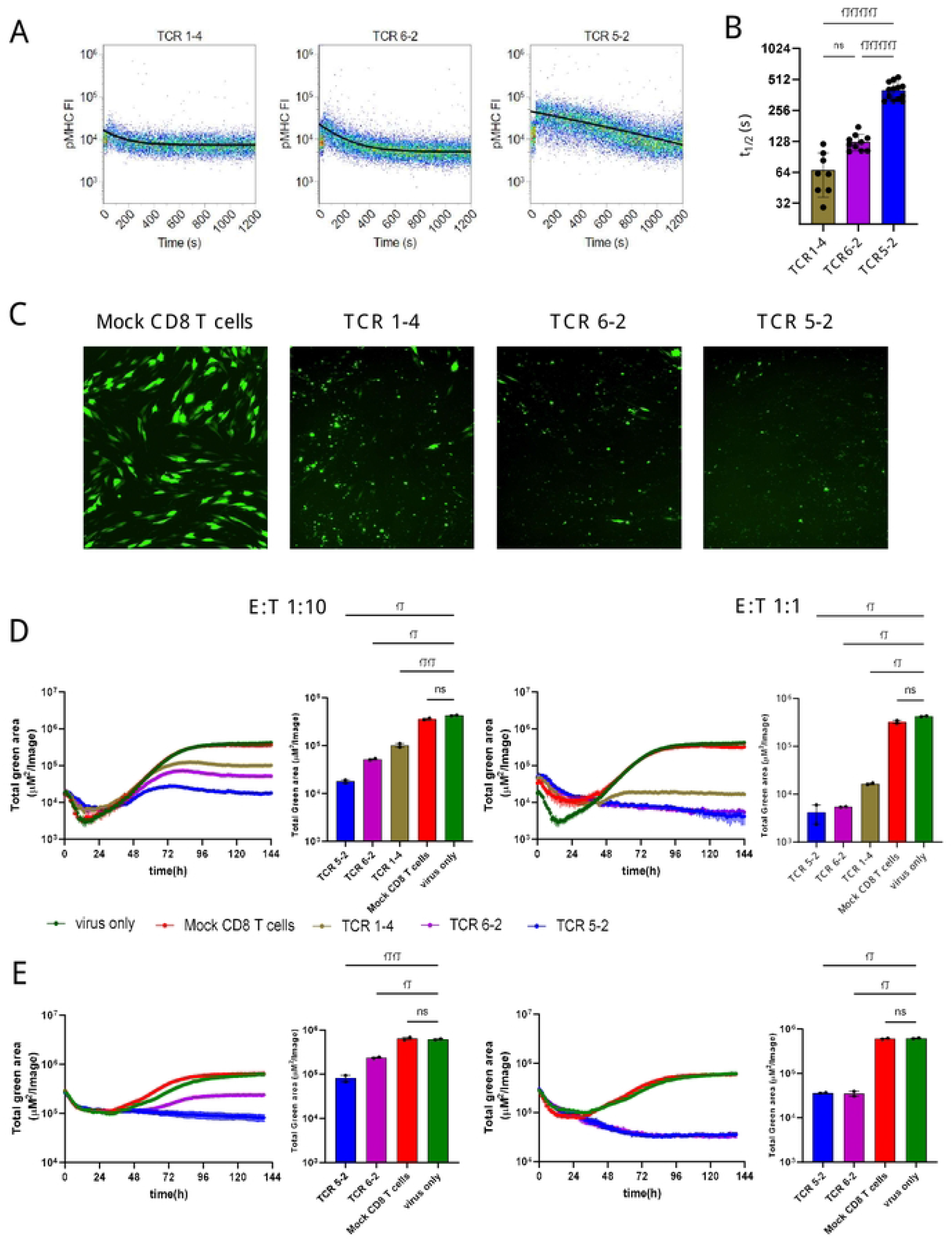
Antiviral activity of transgenic pp65-specific CD8 T cells against TB40^pp65-EGFP^. (A) Representative FACS plots depicting K_off_-rate measurements of TCRs 1-4, 5-2, and 6-2. Lines depict calculated decay of pMHC-monomer fluorescence, indicating dissociation from the TCR. (B) Quantification of half-life times (t1/2) as a measure of pMHC-monomer dissociation kinetics. Each data point corresponds to a single measurement. Means +/-SD are shown. Data are pooled from 3 individual experiments. Statistical analysis was done using Ordinary one-way ANOVA and Tukey’s multiple comparisons test. (C-E) MRC-5 cells were infected with TB40^pp65-EGFP^ at multiplicities of infection (MOI) of 0.1 (C-D) or 1 (E). (C) Representative images of co-culture at 96 hpi showing control of TB40^pp65-EGFP^-infected MRC-5 cells at 1:10 E:T ratio. (D-E) Cells were co-cultured with HLA-A02–restricted pp65-specific CD8 T cells (TCR 1-4 = intermediate avidity, TCR 5-2 and TCR 6-2 = high avidity) at 1:10 (left panels) or 1:1 (right panels) E:T. “Total green area” on the y-axes corresponds to EGFP-pp65 expression, indicating the combined surface of virus infected cells and thus a proxy of viral load over time in ARMATA. This value was plotted at 1h intervals for up to 144 hours post infection. Histograms represent the endpoint value of the total green area for indicated conditions. Each data point corresponds to biological replicates (n=2). Means ± SEM are shown. Statistical analysis was done using Welch ANOVA and Dunnett’s T3 test. ****P < 0.0001, **P < 0.01, *P < 0.05, P > 0.05 not significant (ns).

### Characterization of early antiviral activity of transgenic pp65-specific T cell clones

In co-culture with the HCMV ^pp65-EGFP^ we were able to measure antiviral activity as early as 36-48 hpi, depending on MOI and on E:T ratios. To validate if this was consistent with T-cell activation, we measured cytotoxic effector molecules or cytokines in the supernatants of co-cultures of virus-infected cells and TCR-transgenic T cells or control-infected cells at 36 hpi. We observed significantly elevated levels of IFN-γ, granzymes A and B and FasL in co-cultures of TCR-transgenic cells (Fig. 3A). Notably, the levels of T-cell mediators were higher in co-cultures of T cells with the high-avidity TCR 5-2 than in those with the lower avidity TCR 1-4 (Fig. 3A), consistent with their antiviral activity (Fig. 2). The high levels of cytokines at 36 hpi raised the question why was the antiviral activity detected only after this time point? We considered that pp65 is a tegumental protein, and hence present at high levels immediately upon virus entry into cells. In line with that, the levels of the pp65-fused GFP decreased in the first 12-24 hours before raising again, likely due to the *de novo* transcription and gene expression (Fig.2). This also implied that the signal of preformed and virion-associated pp65-GFP would not be reduced by T cells, thwarting the detection of early antiviral effects. Hence, to monitor earlier events, we used the HCMV expressing mNeon-Green (mNG) in the IE1/3 gene expression context (TB40^IEmNG^), which shows no signal at the time of infection, but a rapid onset by 3-4 hpi (22). While all cells transfected with a pp65-specific TCR reduced the mNG signal, T cells transfected with the high avidity TCR 5-2 depressed the signal more efficiently and showed an earlier onset of activity (Fig. 3B). The effects were more pronounced at a higher E:T ratio, where the differences in virus control by higher and lower avidity TCRs were more pronounced (Fig. 3C).

**Figure 3.**
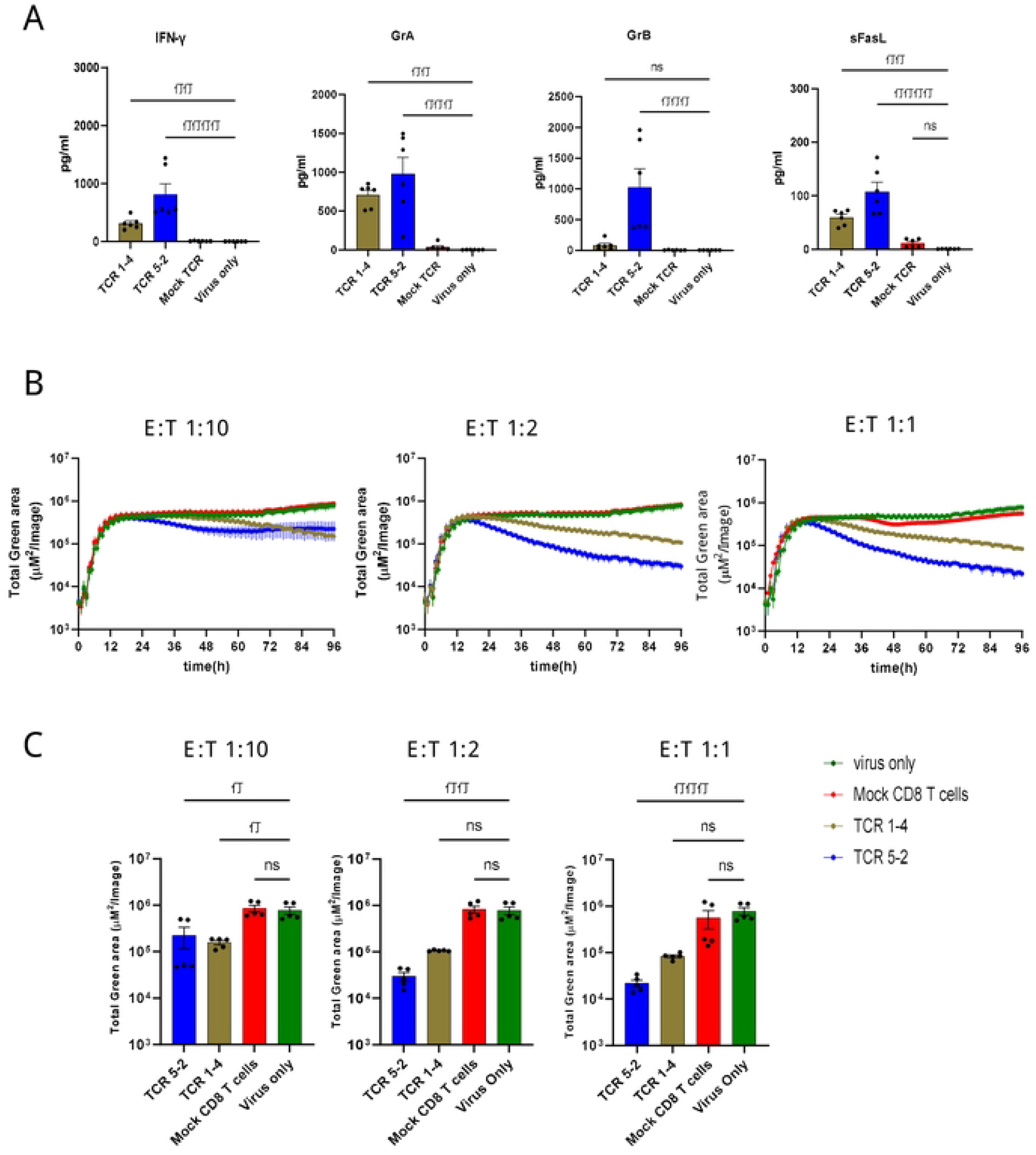
Activity of transgenic pp65-specific CD8 T cells. Cells were infected at MOI of 0.1 in the presence of T-cells transfected with indicated TCRs, or in indicated control conditions and assessed by cytokine concentrations in the supernatant or by ARMATA. (A) Cytokine concentrations in supernatants of 6 independent biological replicates of 1:2 E:T co-cultures at 36 hpi. Results are pooled from three independent experiments, using CD8 T cells, originated from three different PBMC donors. (B) MRC-5 cells were infected with TB40^IEmNG^ at 0.1 MOI and co-cultured with CD8 T cells transfected with indicated TCRs at 1:10, 1:2 and 1:1 E:T. Total Green area indicated on the y-axes corresponds to mNeonGreen-ie1/2 expression indicating viral load plotted for up to 96 hpi in the ARMATA. (C) Representative plots showing endpoint green area. (B, C) Each data point correspond to 5 independent biological replicates pooled from two independent experiments and CD8-T cells isolated from two donors. Histograms depict mean ± SEM. Statistical significance was calculated with Kruskal-Wallis test followed by Dunns post-analysis. *P < 0.05 **P < 0.005, ***P < 0.001 ****P < 0.0001, P > 0.05 not significant (ns)

### Early presentation and recognition of pp65 on MHC class 1 by pp65-specific T cells

To understand how early the antiviral activity of pp65-specific T cells can be identified by ARMATA, we zoomed into the early time points upon infection, within the first 24 hpi (Fig. 4A). For this purpose, co-culture conditions were optimized by using viral inoculum of MOI 1 for robust virus gene expression in target cells. CD8 T cells transgenically expressing the high avidity TCR 5-2 were added to infected cells at E:T ratio of 1:1, 1:2 and 1:10, and dose-dependent effects of T cell preparations were observed at 24 hpi (Fig. 4B). Surprisingly, monitoring at E:T of 1:1 and 1:2 revealed that CD8 T cells inhibited viral gene expression at very early time points, starting around 6 hpi (Figure 4A). In contrast, mNeonGreen-ie1/2 expression started to decline only around 12 hpi when cells were inoculated at 0.1 MOI (Figure 3B), arguing that TB40^IEmNG^ provides an early detection of antiviral activities of CD8 T cells, but that the system is particularly rapid at higher MOI, in single-step growth kinetics conditions. TCR 1-4 also showed a dose-dependent reduction of virus signal in these settings, but the earliest clear differences were spotted only by 10-12 hpi (Fig. S2A). Viral clearance by TCR 1-4 T cells was significant at E:T 1:1, decreasing at 1:2 and not detectable at 1:10 E:T (Fig. S2A, S2B).

**Figure 4.**
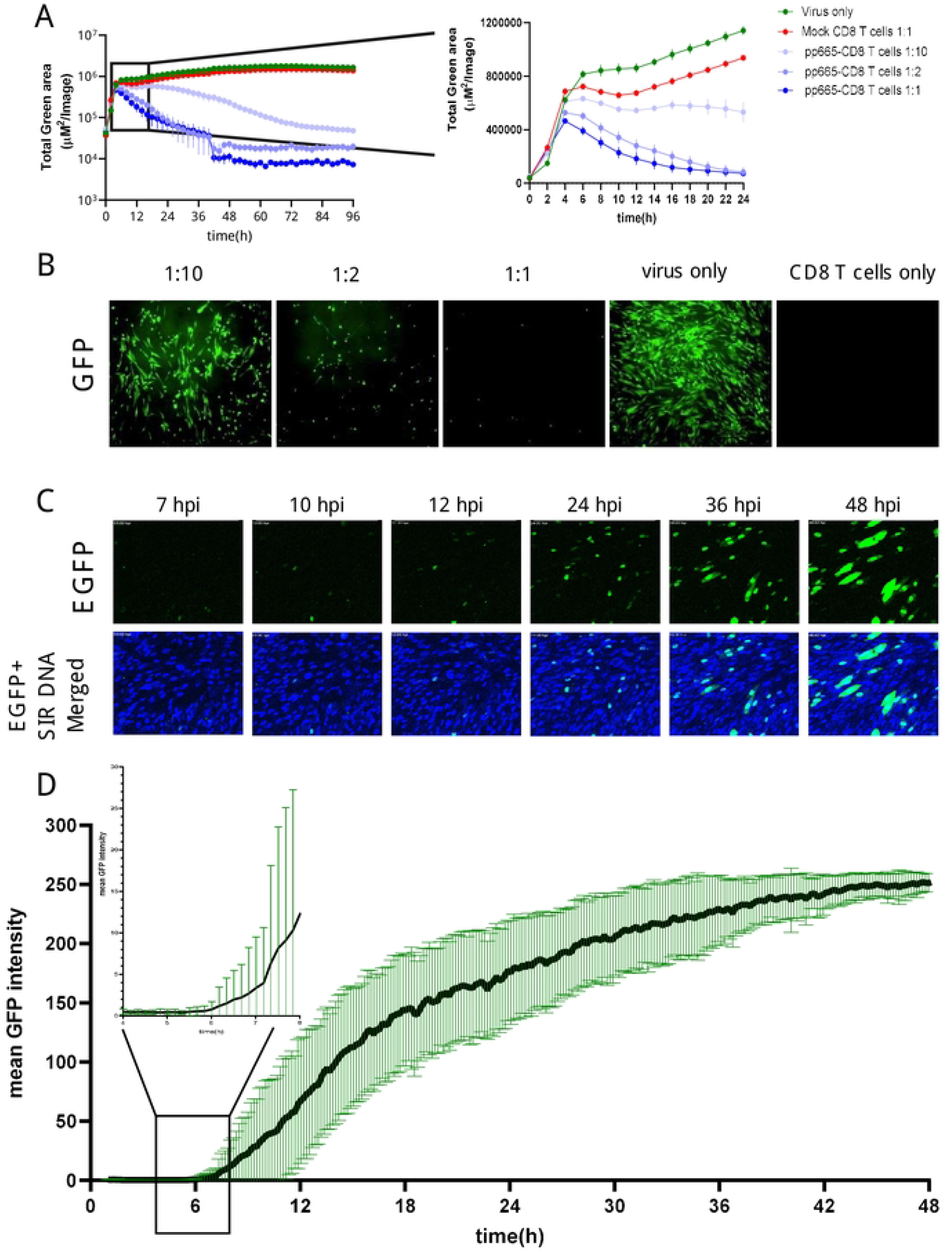
Early expression of pp65 gene and recognition by pp65-specific CD8 T cells. (A) MRC-5 cells were infected with TB40^IEmNG^ at 1 MOI and co-cultured with pp65-specific CD8 T cells transfected with the high-avidity TCR 5-2 at 1:10, 1:2 and 1:1 E:T. Total Green area indicated on the y-axes corresponds to mNeonGreen-ie1/2 expression indicating viral load plotted for up to 96h (Left panel) or 24 hours (Right panel) post infection in ARMATA. Each data point corresponds to the average ± SEM of biological duplicates from two independent experiments. (B) Representative images of co-culture at 24 hpi showing control of TB40^IEmNG^-infected MRC-5 cells at indicated E:T ratios (C) Still frames from live-cell imaging of MRC-5 cells infected for indicated time with TB40^pp65-EGFP^ at 0.1 MOI. Reporter virus expressed EGFP (green) and cells were labelled with Sir DNA (Blue). (D) pp65 gene expression dynamics of TB40^EGFP-pp65^ at the single cell level. MRC-5 cells were infected with TB40^EGFP-pp65^ at MOI 0.1, and subjected to live-cell confocal microscopy acquiring images at 10 min intervals. Data corresponds to average fluorescence profiles of single cells for EGFP-pp65 n= 33]. To visualize the earliest signal rising above the background noise, the signal at 4-8hpi is enlarged in the blow-out square on the left. Data indicate the means ± SD.

In light of these results, we monitored the early expression of the pp65 gene to correlate it to the initial antiviral activity from TCR 5-2 co-cultures. We used purified virus to minimize noise from pp65-GFP molecules in the GFP signal early upon infection and monitored the reporter gene expression in cells infected with TB40^EGFP-pp65^ by confocal live-cell microscopy at high temporal resolution (Figure 4C, 4D), by averaging the GFP signal at single cell levels. Earliest detectable signals started around 6 hpi, which was in line with the antiviral activity of T cells. Taken together, these data suggested that the large expression levels of pp65 at the late stage of the virus cycle are not required for antigenicity, and consequently that pp65 may act as an early gene for immunological purposes.

### Functional activity of pp65-specific T cells in the presence of late gene inhibitor

While pp65 was traditionally considered an early/late viral gene (20), more recent evidence argues that its transcription proceeds even when mRNA translation is inhibited (21), arguing that pp65 may be an immediate-early gene. To distinguish between the dynamics of early and late pp65 expression, we infected MRC-5 cells with TB40^EGFP-pp65^ and blocked DNA replication (and thus by definition, the true late gene expression) by phosphonoacetic acid (PAA) treatment. While the pp65.GFP signal kept increasing in absence of PAA, it flattened by 30 hpi in PAA presence and did not increase further (Fig. 5A). However, the pp65-GFP signal started increasing before 30 hpi and this effect was not repressed by PAA, demonstrating directly that pp65 is not a true late gene (Fig. 5B). This correlated with our data, where pp65-specific CD8 T cells repressed IE gene expression by 6 hpi (Fig. 4A). To test directly if pp65 acts as an early or as a late antigen, we treated the co-cultures of infected cells and TCR 5-2 or TCR 1-4 T cells with PAA, and monitored cytokine responses and viral gene expression. We used HCMV^3F^ for infection, because it expresses the mNG by the IE1/3 promoter and mCherry by the late m48.2 promoter, allowing us to track simultaneously an immediate-early and a late signal (27). Before proceeding with ARMATA in presence of PAA, HCMV^3F^ was tested in its absence (Fig. S3), where TCR transgenic T cells repressed the IE1/3 mNG signal in a manner that matched the effects previously observed with TB40^IEmNG^ (see Fig. 3). Moreover, a clear PAA-induced reduction of IE1/3 and m48.2 signals from HCMV^3F^ was observed in absence of T cells (Fig. S4A) or in the presence of mock-transfected cells (Fig. S3B) starting around 24-36 hpi, in line with previous observations (27). HCMV^3F^ infected cells were then co-cultured in presence of PAA with TCR 1-4 and TCR 5-2 T cells, and the IE1/3 or the m48.2 signals were compared to control co-cultures, where mock-transfected T cells were used. While no late gene expression could be observed in any condition due to PAA treatment (Fig. 5C, right panel, see also Fig. S4), the IE1/3 signal was present in all settings, but reduced in presence of TCR 1-4 and TCR 5-2 (Fig. 5C, left panel). Again, the high avidity TCR 5-2 T cells repressed the signal more efficiently than the TCR 1-4 (Fig. 5C). Finally, we tested the IFN-γ secretion in presence of TCR 5-2 T cells and PAA and observed a clear induction at 24 hpi and an even stronger response at 48 hpi (Fig. 5D). Importantly, no significant difference in INF-γ levels were observed in TCR 5-2 co-cultured CD8 T cell in presence or absence of PAA. In conclusion, pp65 antigenicity was maintained in absence of DNA replication, directly showing that pp65 acts as an early antigen and not a late one.

**Figure 5.**
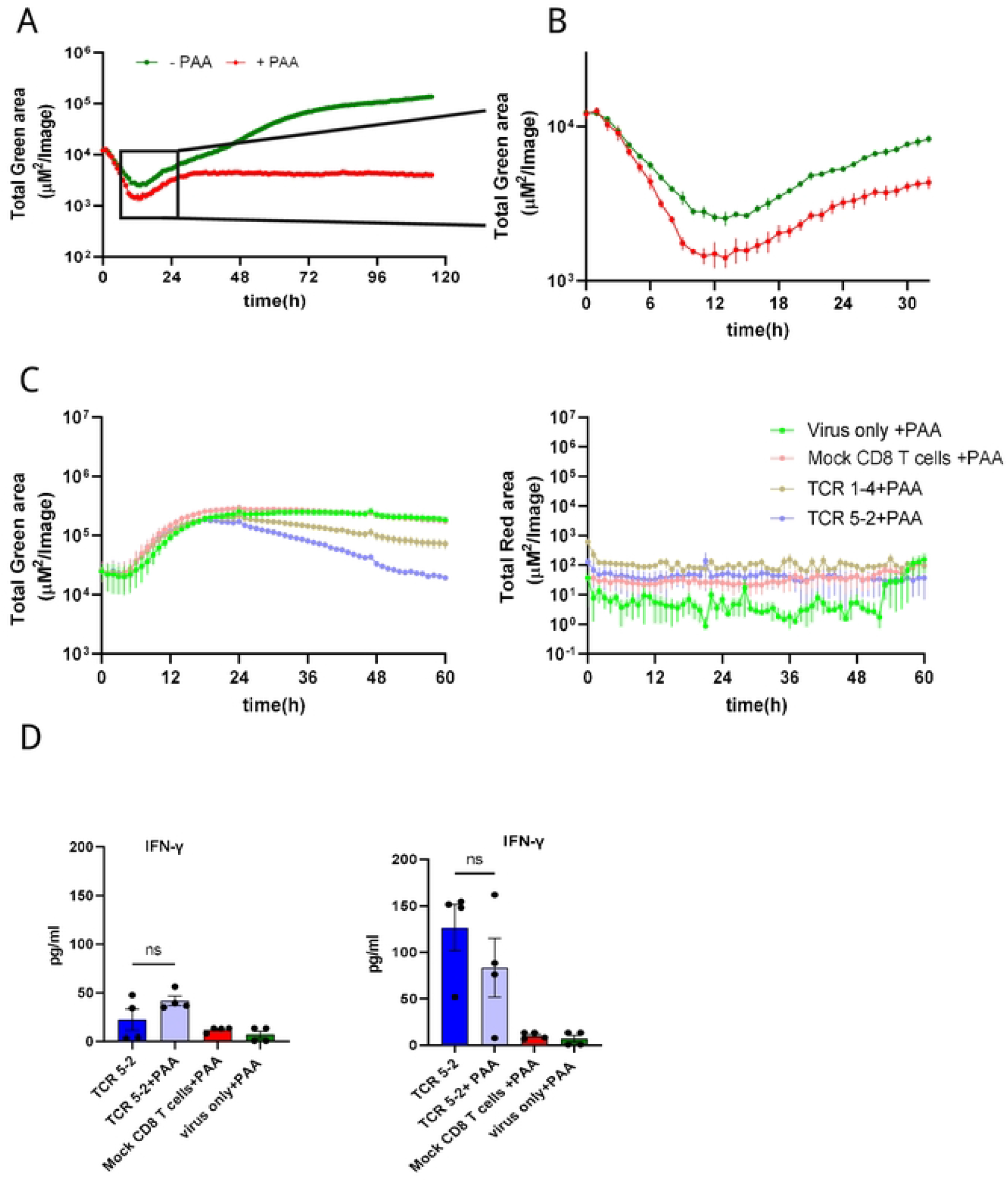
Control of TB40 by different pp65-specific CD8 T cells in the presence/absence of PAA. MRC-5 cells were infected with TB40^pp65-EGFP^ at MOI 0.1 and either left untreated or treated with PAA (100 μg/ml) at the time of infection. Green area on y-axes corresponds to EGFP-pp65 expression indicating viral load plotted in ARMATA for up to 120 hpi (A) or 30 hpi (B). (C) MRC-5 cells were infected with HCMV^3F^ at MOI 0.1 and co-cultured with HLA-A02– restricted pp65-specific CD8 T cells at 1:2 E:T. Cells were treated with PAA (100 μg/ml) at the time of infection. Total Green area shown on y-axes of graphs on the left side corresponds to mNeonGreen-ie1/2 and Total Red area shown on y-axes of graphs on the right side represent UL48A-mCherry expression, indicating total viral loads plotted for up to 60 hpi. For better reference, we show a comparison of the same signals in presence of PAA in infected cells that were co-cultured with CD8 T cells bearing the high-avidity TCR or those with the intermediate-avidity TCR. (D) Cells were infected in same conditions and cytokine concentrations in co-culture supernatants from high-avidity TCR were measured at 24hpi (Left panel) and 48hpi (Right panel). Each data point correspond to independent biological replicates (n=4) from two individual experiments. Histograms with error bars depict mean ± SEM. Statistical significance was calculated via Kruskal-Wallis test followed by Dunns post-analysis. *P < 0.05 **P < 0.005, ***P < 0.001 ****P < 0.0001, P > 0.05 not significant (ns)

## DISCUSSION

While numerous studies have previously addressed the kinetics of pp65 gene expression and its immunodominance, this is to the best of our knowledge the first effort to quantify the antiviral potential of pp65/NLV-specific CD8 T cells and their timing in the context of the virus infection. Importantly, we show that CD8 T cell-mediated virus control in our model system directly correlates to TCR binding avidity highlighting that this approach enables accurate resolution of TCR quality differences. In contrast, assays using peptide-pulsed antigen presenting cell lines were not able to resolve functional differences between those TCRs, at least in the presence of the CD8 co-receptor (24). Furthermore, HCMV is a slow-growing virus, whose replication cycle is typically around 48h in tissue culture. A functional assay to define the antiviral activity of CD8 T cells by flow cytometry of infected cells has been recently described, but the assay requires an incubation of 10-14 days prior to cell harvest (28). Likewise, classical neutralization assays are based on the reduction of viral plaques, which require about a week to form. By measuring early gene expression, we reduced the time required for the identification of antiviral effects to less than 24 hpi, which has significant pragmatic benefits for the rapid identification of effective cell product and may allow the use of our approach in quality controls of cellular products before their administration to a patient.

NLV is a known immunodominant antigen, where CD8 responses accrue in older age groups (29). Similar dynamics of responses were observed in the mouse model of CMV infection (30) and aptly termed memory inflation (31). Memory inflation is related to immediate-early gene expression (32) that cannot be repressed by viral immune evasion genes (33), and which requires processing by the constitutive proteasome (34) in non-hematopoietic cells (35), which are the major sites of virus latency (36). Therefore, it has been proposed that antigenic epitopes that are expressed intermittently during virus latency drive inflationary responses (37) and may serve as optimal targets for protective viral responses (38). Here we demonstrate that the inflationary NLV epitope is a robust target for T cells in the context of the virus infection and that its recognition mediates efficient virus control.

Surprisingly, the antiviral effects could be observed very early, but not immediately upon infection. These results may argue that NLV-specific T cells did not recognize antigenic peptides derived from the tegumental pp65 of the incoming virus, but rather that *de novo* translated products acted as antigens. The timing of the initial virus repression coincided with the earliest detectable signal driven by the endogenous pp65 promoter, arguing that low levels of *de novo* synthesized protein may be processed by the proteasome and presented on HLA molecules much more efficiently than the tegument-associated incoming protein. This may be in line with models where defective ribosomal products (DRIPs) are rapidly recruited to the proteasome for MHC-I presentation (39). However, our data cannot discern if tegumental pp65 failed to act as antigen because of localization (e.g. in the endosomal compartment versus cytosol), due to its accessibility to the proteasome, or due to other factors unrelated to proteasomal processing.

Among the limitations of our work, one has to point out that the effects were measured by using a recombinant CMV lacking the US2-US6 genes, where several among them act as immune-evasive genes limiting the HLA-display of antigens on the cell surface (40). Therefore, it is possible that a recombinant HCMV expressing these genes may be more resistant to CD8 T cells recognizing the NLV epitope. However, we observed antiviral activity at very early stages, before a robust level of immune evasins might limit the delivery of antigen to the cell surface. Since immune evasins preferentially affect cell surface display of recently generated peptide presentation complexes, and are poor at downregulating those that already exist (41), and since we compared antiviral efficiencies of various transgenically expressed TCRs, we consider our data representative of *in vivo* effects that such cells may exert. However, additional work with recombinant viruses expressing immune evasins is required, especially for viral antigens that are expressed later in the virus replication cycle. Another limitation is that we explored the effect of only one antigenic epitope. Future work based on this model should compare various viral antigens and serve to identify those that result in the best control of the virus growth. This study paves the way towards such approaches, providing the proof of principle that dynamic monitoring of virus signals allows a rapid identification of cell populations with optimal antiviral activity.

## Material and Methods

### Cell Lines

MRC-5 (ATCC Cat# CCL-171) cells were purchased from American Type Culture Collection. Cells were cultured in MEM supplemented with 10% FCS, 20 nM glutamine and 1 mM sodium pyruvate and maintained at 37°C, 5% CO2, and 100% air humidity.

### Virus generation and stock production

Generation of the reporter TB40^IEmNG^ has been described in detail previously (42). Briefly, mNeonGreen gene linked by the P2A peptide was inserted before the start codon of UL122/123 exon 2. The same strategy was used in HCMV^3F^, where additionally the mCherry was linked by a T2A peptide to the UL48a, as described (23). TB40^EGFP-pp65^ was generated by fusing the open reading frame of EGFP with the UL83 gene resulting in EGFP expression under the UL83 promoter (43).

Virus stocks were generated on MRC-5 cells. Cell supernatants and lysates were centrifuged (500 × g, 10 min) to remove debris, followed by viral particle concentration via centrifugation at 24,000 × g, for 3.5 h at 4°C. Pellets were homogenized by douncing and subjected to ultracentrifugation for 90 minutes through a 15% sucrose-cushion for additional purification. Virus stocks were titrated on MRC-5 cells as described before (42)

### Cell Culture and Viral Infection

Cells were seeded into 24-well plates at 80,000 cells/well or in 96-well plates at 20,000 cells/well. Infection with HCMV was done by diluting the virus in cell culture medium to indicated MOI (where MOI 1 means 1 PFU/cell and MOI 0.1 means 1 PFU per 10 cells), followed by a centrifugation of cells for 10 min at room temperature at 800 x g.

### Human Peripheral blood mononuclear cells (PBMC) Isolation

Leukoreduction system (LRS) chambers for PBMC isolation from healthy blood donors (CMV and HLA-A02-negative) were supplied by the Institute for Transfusion Medicine, MHH under the ethical permit 2519-2014 from MHH Ethics commission. PBMCs for retroviral transduction were isolated from LRS chambers by standard density gradient centrifugation procedure.

### Retroviral transduction of Human PBMCs

TCR plasmids with respective sequences were generated at TUM and retroviral transduction of human CD8 T cells was performed at HZI as previously described (13). Briefly, RD114 cells were seeded at 2 × 10^5^ cells/well in a 6-well plate with DMEM (Gibco, USA) supplemented with 10% FCS (Sigma-Aldrich, USA), L-glutamine (Gibco, USA), and transfected 2-3 days later with pMP71 expression vector containing the TCR sequence. Transfection was performed via calcium phosphate precipitation with 18 μg of the vector DNA in 15 μL of 3.31 M CaCl_2_ mixed with 150 μL of the transfection buffer (containing 274 mM NaCl, 9.9 mM KCl, 3.5 mM Na2HPO4 and 41.9 mM HEPES) and incubated for 30 min at RT. The mix was added dropwise to RD114 cells and incubated for 6 h at 37°C, 5% CO2 followed by a complete medium exchange. Virus supernatant was harvested three days later and coated on retronectin (TaKaRa)-treated plates (incubated overnight at 4°C previously) via centrifugation at 2000 g, for 2 h at 32°C.

For PBMCs stimulation, 24-well plates were coated with 250μL PBS/well containing 5 μg/mL aCD3 + 0,05 μg/mL aCD28 and incubated overnight at 4°C. Coated plates were blocked with 2% BSA in PBS for 20 min at 37°C, then washed twice with PBS. 1×10^6^ PBMCs/well were seeded in RPMI medium + 10% FCS + 300 IU/mL human IL-2 and incubated for 2 days.

Activated PBMCs were spinoculated onto the virus-coated plates (1000 g, 10 min at 32°C). Transduced cells were kept in RPMI medium +10% FCS + 180IU/mL human IL-2 for 5-7 days. At the end of transduction, efficiency was validated by flow cytometry staining of CD8+-mTCR+ (murine constant TCR-β chain) cells via flow-cytometry.

### Transduced CD8 T cell selection with Manual MACS columns

Fresh/cryopreserved transduced PBMCs were rested and then processed to isolate CD8+/mTCR+ population. Briefly, cells were collected and centrifuged at 500 x g for 5 min at 4°C and CD8 T cells were isolated by magnetic sorting according to the manufacturer’s protocol (Miltenyi Biotec, Germany). To select transduced CD8 T cells, a second round of selection based on the mTCR-β chain was performed. Cells were incubated with PE-anti-mTCR-β antibody for 20 min at 4°C, followed by centrifugation and washing. Cells were magnetically labelled with anti PE-microbeads (Miltenyi Biotec, Germany) and positive cells were collected from the column. Purity of isolated T cells was determined via flow cytometry.

### K_off_-rate measurement

K_off_ rates were determined as dissociation of reversible pMHC-Streptamers upon addition of D-Biotin in a flow cytometry-based assay (CyAn ADP Lx 9 color flow cytometer, Beckman Coulter) as described previously (44). In brief, pMHC molecules were multimerized with Strep-Tactin APC (IBA lifesciences) according to the manufacturer’s instructions and incubated with 5×10^6^ TCR-transgenic T cells for 45 min. After 25 min, cells were additionally stained with CD8α PE (clone OKT8, eBioscience). For live/dead discrimination, cells were incubated with propidium iodide (Invitrogen). 1×10^5^ - 1×10^6^ pre-cooled cells were analyzed by flow cytometry. After 30 s, D-Biotin was added to a final concentration of 1 mM while analysis continued. Dissociation kinetics were analyzed using FlowJo (FlowJo, LLC) and GraphPad Prism software (GraphPad Software).

### Microscopy and Image Analysis

Time-lapse imaging was performed either in Incucyte-S3 devices (Sartorius) or in a ZEISS LSM 980 confocal laser-scanning microscope (Carl Zeiss Microscopy). Images from Incucyte-S3 were analyzed by Incucyte™ GUI Software (versions 2019B REV1 or 2021B), while time series image stacks from the confocal microscope were analyzed with Fiji (ImageJ). Single cell tracking was performed by a manual in-house Fiji macro. Infected cells were counted for mean GFP signal intensity from each frame manually. Data was plotted using GraphPad Prism.

### Assay for Rapid Measurement of Antiviral T-cell Activity

HLA-A02-positive MRC-5 cells were infected with reporter HCMV by centrifugigation at 800 x g for 10’ at RT. Immediately upon infection, we added pp65-specific CD8 T cells in CTS OpTimizer T Cell Expansion Serum Free Medium medium (Gibco), supplemented with 5% CTS™ Immune Cell SR (Gibco) and 20 U/mL human IL-2 (PeproTech). Co-cultures were monitored by Incucyte™ imaging system using the green and red fluorescent channel for reporter gene expression. Images were acquired at regular intervals and analyzed by the Incucyte™ GUI Software.

### Cytokine analysis from co-culture

Cytokine analysis from co-culture supernatants was performed with Human CD8/NK Panel (13-plex) Cat#741065 (BioLegend) following the manufacturer’s protocol.

### Statistical Analyses

Where relevant, non-parametric statistical analysis was used. Kruskal-Wallis test followed by Dunn’s postanalysis was performed with Graphpad prism 9.0 to quantify the statistical significance in multiple comparisons. P values < 0.05 were considered significant (*P < 0.05, **P < 0.01, ***P < 0.001, ****P < 0.0001); P > 0.05 not significant (ns).

## Supplementary Figures

**Supplementary figure 1. Flow cytometric validation of transgenic CD8+mTCR+ cell purification efficiency**

CD8 T cells were transfected with retroviruses expressing the indicated TCR and grown for 3-7 days. Transduction efficiency was validated by Flow cytometry staining of CD8^+^-mTCR^+^ cells (upper panel) and the same was performed upon MACS-sorting of mTCR^+^ CD8 T cells (lower panel) to control the purity of T-cells used in co-culture experiments. Representative purities are shown, no cultures with <80% of mTCR^+^ cells have been used in any experiment.

**Supplementary Figure 2. Early expression of pp65 gene and recognition by pp65-specific CD8 T cells**

MRC-5 cells were infected with TB40^IEmNG^ at a MOI of 1 and co-cultured with pp65-specific CD8 T cells transfected with the Intermediate-avidity TCR 1-4 at 1:10, 1:2 and 1:1 E:T. Total Green area indicated on the y-axes corresponds to mNeonGreen-ie1/2 expression, indicating viral loads plotted for up to 96h (Left panel) or 24 hours (Right panel) post infection in ARMATA. Each data point correspond to the average ± SEM of biological duplicates from two independent experiments. (B) Representative images of co-culture at 24 hpi showing control of TB40^IEmNG^-infected MRC-5 cells at indicated E:T ratios of Intermediate-avidity CD8 T cells.

**Supplementary Figure 3. Antiviral activity of transgenic pp65-specific CD8 T cells against HCMV**^**3F**^

(A) MRC-5 cells were infected with TB40/HCMV^3F^ at MOI of 0.1 and co-cultured with CD8 T cells transfected with indicated pp65-specific TCRs at 1:10, 1:2 and 1:1 E:T. Total Green area indicated on the y-axes corresponds to mNeonGreen-ie1/2 expression indicating viral loads plotted for up to 120 hpi in ARMATA. (B) Representative plots showing endpoint green area. Each symbol corresponds to a biological replicate at indicated E:T ratios, pooled from ≥3 independent experiments. Histograms depict means ± SEM. Statistical analysis was done using Welch ANOVA and Dunnett’s T3 test. **P < 0.01, ***P < 0.001 ****P < 0.0001, P > 0.05 not significant (ns)

**Supplementary Figure 4. Immediate-early and late gene expression dynamics of TB40**^**3F**^ **reporter genes in the presence or absence of PAA**.

MRC-5 cells were infected with HCMV^3F^ at MOI 0.1 and were either left untreated or treated with PAA (100 μg/ml) at the time of infection. Total Green area shown on y-axes of graphs on the left side corresponds to mNeonGreen-ie1/2 while Total Red area shown on y-axes of graphs on the right side represent SCP-mCherry expression, indicating viral loads in distinct stages of the lytic replication cycle plotted for up to 60 hpi. Each data point correspond to independent biological replicates (n=4) from two individual experiments. Data with error bars depict mean ± SEM.

## Notes

### Competing Interest Statement

HZI holds a patent on MCMV-3F, one of the viruses used in this study. LCS is one of the inventors listed on the patent.

